# Computational Analysis of ELOVL6 Structure and Inhibition for Rational Drug Design

**DOI:** 10.1101/2025.08.23.671927

**Authors:** Markel G. Ibarluzea, Rafael Ramis, Martin Fuentetaja Leza, Francisco J Gil-Bea, Gorka Gerenu, Adolfo López De Munain, Jesus M. Aizpurua, José I. Miranda, Aitor Bergara, Aritz Leonardo

## Abstract

ELOVL6 is a key enzyme in long-chain fatty acid elongation, catalyzing the conversion of C16 fatty acids into C18 fatty acids. While its role in lipid metabolism is well established, recent studies have linked ELOVL6 to metabolic and neurodegenerative diseases, making it an attractive therapeutic target. However, the absence of a resolved crystal structure and limited mechanistic understanding of its inhibition pose significant challenges for drug discovery.

In this study, we employ a multi-tiered computational approach, including structure prediction, molecular dynamics (MD) simulations, and free energy calculations, to investigate the structural basis of ELOVL6 function and inhibition. We identify the most thermodynamically favorable substrate binding pathway and characterize key conformational changes associated with ligand binding. By analyzing potential inhibitor binding pockets, we determine that known inhibitors preferentially target the active site, and we validate their binding affinities against experimental data. Additionally, by comparing ELOVL6 with homologous elongases, we pinpoint potentially key amino acid residues responsible for selectivity, providing insights that could guide structure-based drug design.

Our findings establish a mechanistic framework for rational inhibitor development, offering a foundation for future efforts in optimizing ELOVL6-targeting therapeutics.

## INTRODUCTION

The ELOVL gene family encodes a group of microsomal enzymes responsible for the elongation of long-chain fatty acids (FAs). These enzymes catalyze the rate-limiting step in fatty acid elongation by adding two-carbon units to acyl-CoA precursors through a series of condensation reactions. ELOVL proteins play a crucial role in lipid metabolism, contributing to the biosynthesis of essential FAs, which are involved in lipid signaling, energy metabolism, and serve as key components of membrane lipids.

To date, seven members of the ELOVL protein family have been identified, each with distinct substrate specificities and roles in lipid metabolism. ELOVL6, a member of this family, has garnered significant attention due to its link to obesity-related malignancies [1, 2, 3]. ELOVL6 is predominantly expressed in the liver and adipose tissue and is responsible for the elongation of C16 FAs into C18 FAs, specifically catalyzing the conversion of palmitoyl-CoA into stearoyl-CoA and palmitoleoyl-CoA into cis-vaccenoyl-CoA.

Most research on ELOVL6 has focused on its role in cardiovascular and metabolic-related malignancies, such as type 2 diabetes, nonalcoholic steatohepatitis, and atherosclerosis [2, 3, 4]; however, more recently, interest in this enzyme has resurged due to its potential in the treatment of inflammatory, neurological diseases, as well as cancer [5]. For instance, ELOVL6 has been found to be upregulated in immune cells involved in myelin degradation in multiple sclerosis hampering remyelination [6], or dramatically upregulated in the lethal pancreatic ductal adenocarcinoma, hepatocellular carcinoma, acute myeloid leukemia, lung squamous cell carcinoma, and glioblastoma multiforme during tumour progression, while its inhibition suppresses tumour growth [7, 8, 9, 10, 11]. This diverse involvement in pathological processes, spanning from metabolic to nervous system disorders and cancer, underscores the need for pharmacological tools to modulate ELOVL6 activity.

Previous studies [12, 13, 14] have reported highly potent ELOVL6 inhibitor molecules. These inhibitors consisted of two main scaffolds, with various derivatives synthesized and tested by modifying substituents at different R-groups. Several derivatives exhibited strong inhibition, with IC_50_ values below 100 nM, and from these tested molecules, one lead compound from each scaffold was selected (Figure 1), referred to as compounds A and B. These compounds were further tested against ELOVL family members 1, 2, 3, and 5 to assess their selectivity. The results demonstrated that both lead molecules exhibited a strong preference for ELOVL6, with compound A showing approximately 38-fold selectivity and compound B displaying around 7-fold selectivity over ELOVL3, the most relevant homolog when considering ELOVL6 selectivity.

**Figure 1.**
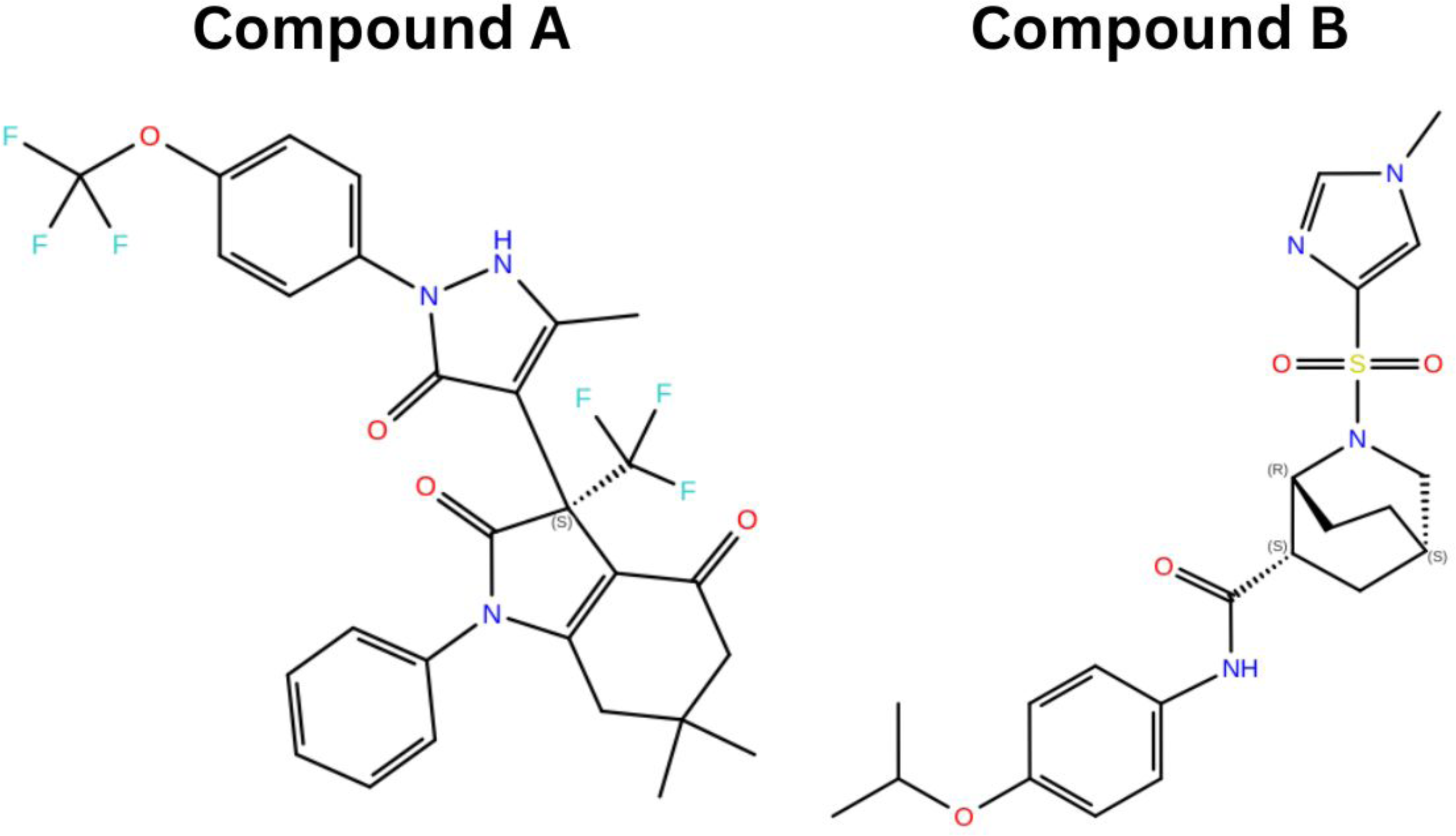
Lead compounds selected from two congenertic series of ELOVL6 inhibitors.

Despite their initial effectiveness, these compounds have failed to reach clinical endpoints. This suggests potential pharmacological limitations or unresolved toxicity concerns that have impeded their progression into clinical trials. Therefore, designing alternative inhibitors with optimal pharmacokinetic properties remains necessary. However, no crystal structure of ELOVL6, in either its apo or holo conformation, has been published to date, making the precise mechanism of action of ELOVL6 and its inhibitors poorly understood. Among the ELOVL family members, ELOVL7 is the only one for which a crystal structure, in complex with 3-keto eicosanoyl (C20)-CoA, is available (PDB ID: 6Y7F). While ELOVL6 and ELOVL7 elongate FAs of different chain lengths and thus likely exhibit structural differences, their high sequence homology (30.45%) suggests that the ELOVL7 structure can provide meaningful insights into ELOVL6’s structure and mechanism of action. However, since this structure was not crystallized with an inhibitor, it provides limited information regarding inhibition mechanisms and binding pockets in ELOVL proteins, particularly ELOVL6. These knowledge gaps significantly hinder the rational design of new inhibitors with high affinity, selectivity, and favorable pharmacological properties for clinical applications.

In this work, we employed multiple computational techniques, including structure prediction algorithms, molecular docking, molecular dynamics simulations, and alchemical free energy calculations, to develop structural models for the binding mechanism of ELOVL6 inhibitors. In a retrospective evaluation, we demonstrated that these models successfully identified a high percentage of known inhibitors from a virtual screening library primarily composed of decoy molecules. Additionally, they accurately ranked different ligands within a congeneric series by affinity, proving their potential to aid in the *in silico* lead optimization process.

## RESULTS

### Substrate binding mechanism analysis

The available crystal structure of ELOVL7 provides valuable insight into the bound conformation of acyl-CoA substrates in elongase proteins. However, the mechanistic details of ELOVL function remain incompletely understood. A two-step ping-pong mechanism has been proposed as the most consistent with available structural data to explain the acyl chain elongation process [15]. For ELOVL6, the first step in this mechanism involves the initial insertion of a palmitoyl-CoA substrate into the active site, where the enzyme facilitates the cleavage of the CoA moiety, leaving behind the acyl chain as an enzyme-bound intermediate. Subsequently, malonyl-CoA enters the active site and undergoes decarboxylation, generating a reactive carbon which attacks the enzyme-bound acyl intermediate, forming a new carbon-carbon bond, thereby extending the fatty acid chain by two carbons to produce stearoyl-CoA.

Despite this proposed framework, key aspects of the elongation process remain unresolved. Notably, the substrate insertion pathway into ELOVL6 is unknown, limiting our ability to rationalize potential mechanisms for its inhibition via ligand binding. One plausible binding pathway involves sequential insertion, where the carbon chain enters the binding site first, followed by the CoA moiety. However, given the narrow dimensions of the binding pocket and the substrate’s inherent flexibility, this process may present a significant entropic bottleneck. Alternatively, binding could proceed via a single-step lateral insertion through the gap between the H4 and H7 helices of ELOVL6. Unlike other helices, H4 and H7 are not connected by a linker chain, potentially allowing sufficient conformational flexibility to widen the gap and accommodate substrate entry.

To determine the minimum free energy pathway for palmitoyl-CoA binding to ELOVL6, we computed the potential of mean force (PMF) for both hypothesized mechanisms. AlphaFold2 [16] was used to generate structural models of ELOVL6, which were subsequently aligned to the ELOVL7 crystal structure (Figure 2A). The structural comparison revealed high similarity, with ELOVL7 being slightly longer to accommodate extended acyl chains. To assess the stability of the generated structure, we conducted molecular dynamics simulations at 300K, 400K, and 500K for both the AlphaFold-generated ELOVL6 model and the ELOVL7 crystal structure. The RMSD and secondary structure preservation remained comparable between the two structures across all temperatures (Figure 1 SI). A palmitoyl-CoA molecule was positioned in the binding site of the ELOVL6 model by superimposing it with the acyl-CoA observed in ELOVL7s binding pocket, followed by energy minimization using Prime [17]. To ensure proper system equilibration, a 20 ns molecular dynamics (MD) simulation was conducted.

**Figure 2.**
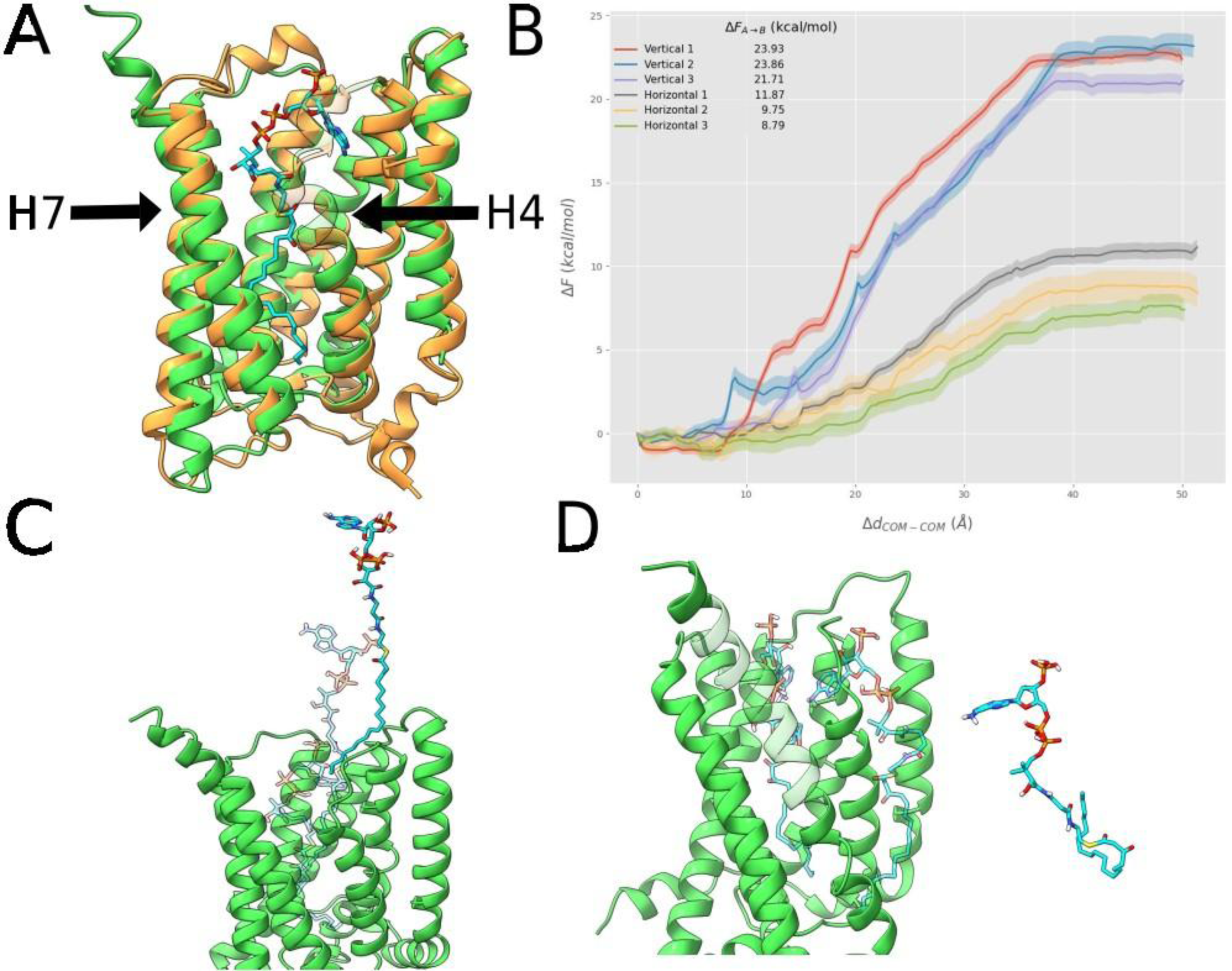
AlphaFold generated ELOVL6 structure, superimposed with crystal structure of ELOVL7 (PDBID: 6Y7F), with a palmitoyl-CoA substrate in the binding pocket, modeled based on the substrate co-crystallized with ELOVL7. (B) PMF profiles associated with each pathway, with 3 replicas per pathway. (C) Structural representations of the hypothesized lateral insertion binding mechanisms through the gap between H4-H7 helices (D) Structural representations of the hypothesized sequential insertion mechanisms, with initial insertion of the carbon chain into the occluded portion of ELOVL6, followed by CoA binding at the active site.

Starting from equilibrated configurations extracted from the last 10 ns of conventional MD, 10 steered MD simulations were performed to generate initial configurations along both hypothesized binding pathways. In these simulations, a harmonic potential with a force constant of 1000 kJ/(mol nm^2^) was applied between the centers of mass of the substrate and helices H2, H3, and H5, pulling at a constant velocity of 0.01 nm/ps over 500 ps. For the sequential insertion pathway, the force was directed towards the active site opening, while for the lateral pathway, it was applied orthogonally, towards the gap between H4 and H7 (Figure 2B). The three trajectories with the lowest accumulated work for each pathway were selected as starting configurations for umbrella sampling simulations.

For each pathway, the umbrella sampling simulations employed a minimum of 25 windows, spaced at 2 Å intervals along the reaction coordinate. Additional windows were introduced where poor overlap between adjacent histograms was observed. Each window was simulated for 75 ns following a 10 ns equilibration period. The PMF was subsequently obtained using the WHAM estimator implemented in GROMACS 2024 [18].

The resulting PMF profiles indicate that the energetic barriers for the lateral pathway are significantly lower than those for sequential insertion, with an average difference of ∼13 kcal/mol across three independent replicas (Figure 2C). This suggests that lateral insertion represents the most thermodynamically favorable binding pathway for ELOVL6. Notably, all trajectories along the lateral binding route exhibited significant widening of the gap between H4 and H7, primarily through displacement of H4, facilitating substrate entry. These observations suggest that substantial conformational rearrangements accompany palmitoyl-CoA binding. These results suggest that targeting this process, either by increasing the energetic cost of H4 displacement or by designing ligands that stabilize alternative conformational states, may represent viable strategies for inhibiting ELOVL6 activity.

### Binding pocket identification

Although multiple potent and selective inhibitors of ELOVL6 have been reported, the lack of a crystal structure for ELOVL6 in either apo or holo conformation leaves the mechanism of action of these molecules poorly understood. Computational studies have previously been used to identify potential binding modes of known inhibitors of ELOVL1 [19], another member of the elongase family. However, the apo structure of ELOVL6 features a binding site volume of approximately 80 Å³ at the location of the suggested ELOVL1 binding site, while some of the larger ligands studied here have volumes around 350 Å³. This discrepancy suggests that significant conformational rearrangements would be required for the binding site to accommodate these inhibitors.

To explore alternative binding pockets in ELOVL6, we conducted mixed solvent molecular dynamics (MD) simulations. In these simulations, ELOVL6 was immersed in a solution of water mixed with small probe molecules with diverse physicochemical properties at a concentration of 2M, following the methodology outlined in [20]. The probe molecules included isopropanol, acetamide, acetate, isopropylamine, and benzene, chosen for their ability to form varied interactions with the target and for their frequent occurrence as fragments in drug molecules. Simulations were performed in five replicas of ∼250 ns each. The resulting trajectories were analyzed by constructing a spatial grid around ELOVL6 and applying the inverse Boltzmann relation to estimate the binding free energy of each probe molecule at each grid point. The values of the individual spatial grids of each simulation were averaged.

Analysis of the probe molecules density on the grid revealed regions of high binding affinity, which could serve as potential binding pockets for inhibitors. Based on these high-affinity volumes able to fit compounds of the size considered in this study, and our understanding of ELOVL6’s mechanism of action, we hypothesized three potential binding pockets (Figures 3A and 3B). The first potential pocket lies in the gap between helices H4 and H7, where ligand binding could block the insertion of FA substrates into the active site. The second pocket is at the edge of the occluded end of ELOVL6, where a ligand binding allosterically could alter the conformation of the substrate binding site, reducing the affinity of either of the two substrates involved in the elongation mechanism for ELOVL6. The third hypothesized pocket is directly at the active site, which exhibited the highest probe molecule density; here, ligand binding would directly interfere with CoA interaction with the active site.

**Figure 3.**
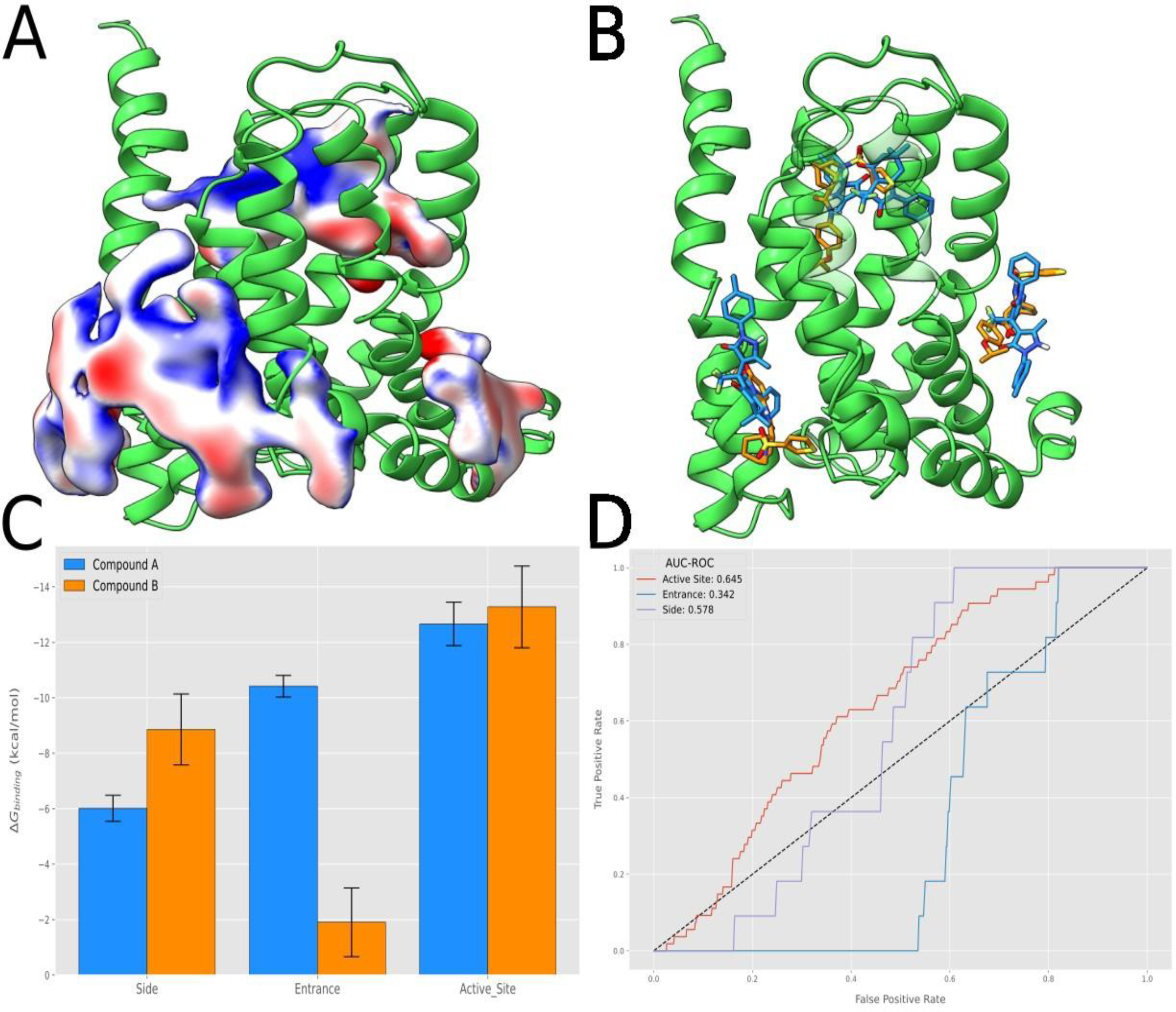
(A) Density maps of probe molecule affinities for ELOVL6, with blue surface representing low densities and red representing high densities. Density below a threshold was removed and remaining densities with insufficient volume to fit the active compounds are not shown for clarity. (B) Binding poses of compounds A and B, obtained from docking these molecules to the three proposed binding sites. (C) Binding free energies of compounds A and B from ABFE calculations for each of the three binding sites. (D) ROC curves of retrospective virtual screenings with a library of decoy molecules from the DUD-E database and experimentally validated active molecules.

To test these hypotheses, we docked two known inhibitors, compounds A and B, into each of the three potential binding pockets. Absolute binding free energy (ABFE) calculations were performed to estimate their affinities using the FEP code in GROMACS 2024, following the procedure outlined in [21]. For two of the three binding pockets, where a significant portion of the ligand is membrane-exposed, a lipid bilayer was included to mimic the composition of endoplasmic reticulum (ER) membranes. The bilayer consisted of 90% POPC and 10% cholesterol, reflecting the lower cholesterol content characteristic of typical ER membranes [22]. The ABFE simulations clearly indicate that the active site binding pocket has significantly higher binding affinities for both compounds, compared to the other two hypothesized binding pockets. The free energy estimates at the active sites were −12.66 (+/−0.78) kcal/mol and −13.27 (+/− 1.47) kcal/mol for compound A and B, respectively, in line with experimentally measured IC_50_ values of 8.9nM and 221nM, which assuming competitive binding and using the Cheng-Prusoff equation, correspond to free energy ranges between −11.7 to −16.4 kcal/mol and −9.8 to −14.5kcal/mol for compounds A and B, using the experimental substrate concentration of 40μM and the K_m_ values of 4nM for palmitoyl-CoA and 11.1μM for malonyl-CoA, as reported in [23]. The binding free energies obtained for the other two models, however, fall well short of the expected values given the known experimental measurements (Figure 3C).

To further benchmark the suitability of these binding pockets, we conducted a set of retrospective virtual screening campaigns for each of the binding pockets, and evaluated the ability of these binding pocket models to differentiate experimentally validated active molecules from a library of decoy molecules with similar molecular properties. The active molecules were obtained from the aforementioned ELOVL6 inhibitor studies [12, 13, 14], by classifying all tested molecules with IC_50_s below 1μM as active, whereas molecules with higher IC_50_s were classified as inactive. The inactive molecule class was augmented with decoy molecules obtained from the DUD-E database [24], resulting in a final ligand library distribution of 1.59% actives (54 molecules), and 98.4% inactives (3343 molecules).

The retrospective analysis aligned with our ABFE calculations, indicating that two binding pocket models—the H4-H7 gap and the allosteric site—failed to enrich for actives, with the H4-H7 gap model performing no better than random selection, and the allosteric site model marginally surpassing it. In contrast, the active site model showed some predictive power, with an AUC-ROC of 0.645 (Figure 3D). However, its performance was inconsistent: for the first approximately 20% of scored molecules, it performs close to random, as reflected by the initial close tracking of the diagonal line by the ROC curve. Visual inspection suggests that the AlphaFold-predicted pocket conformation is too rigid to accommodate larger active molecules and exhibits a bias favoring derivatives of compound A over those of compound B. These findings highlight the need for conformational refinements to improve the predictive utility of this binding pocket model.

### Binding pose identification

Based on the promising results from the active site binding pocket model, we sought to refine this pocket to better reflect the true holo conformation of the protein. To determine the optimal binding poses of known inhibitors from series 1 and 2 (Figure 4A), we implemented a multi-stage protocol (Figure 2 SI) involving the identification and validation of multiple candidate poses. Unlike the analyses in previous sections, which relied solely on the top AlphaFold2 structure, in this case we considered all five AlphaFold2 models. Despite their overall similarity, some relevant differences between these structures can be observed in their modeling of the binding pocket.

**Figure 4.**
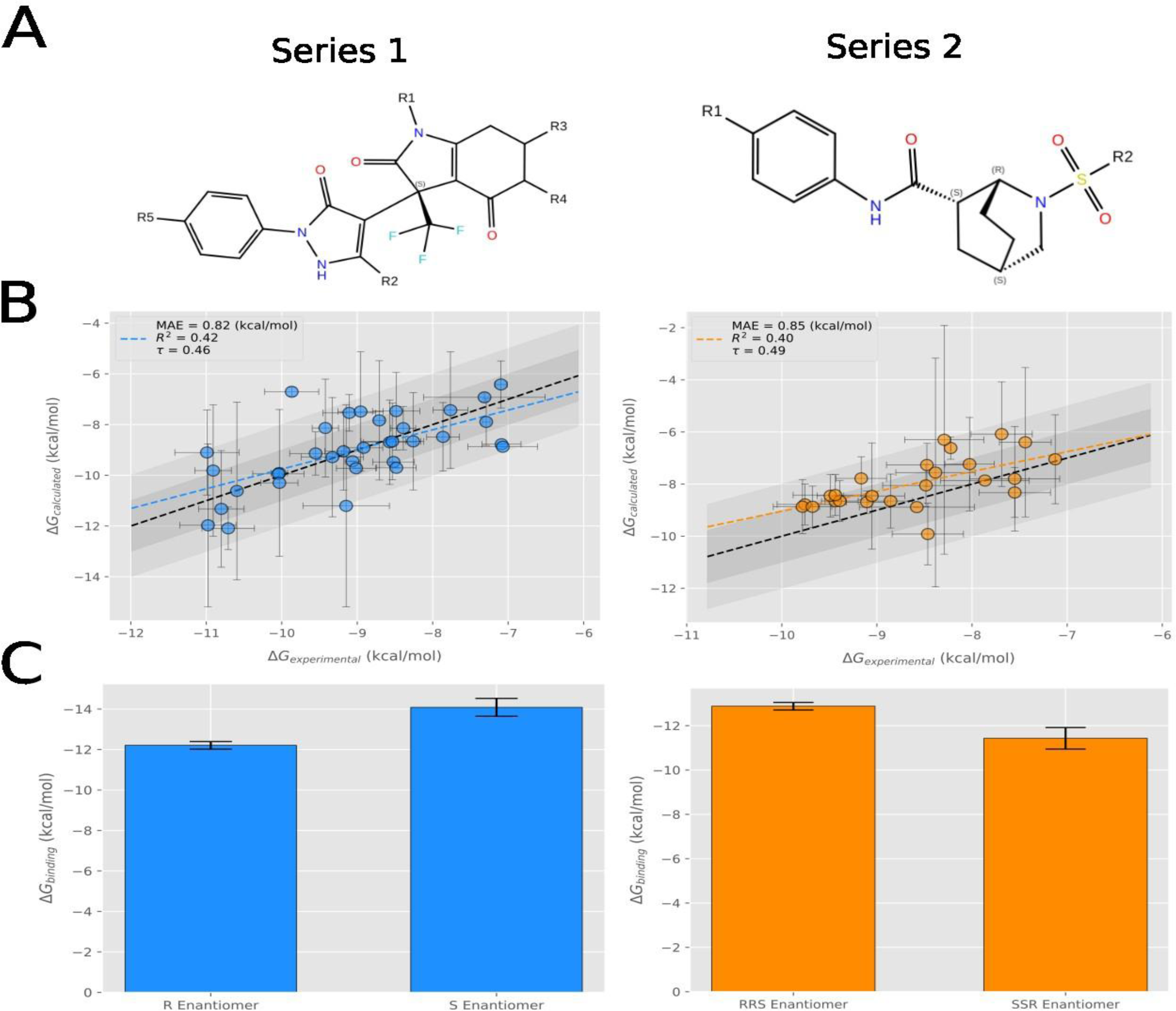
(A) Chemical structures of congeneric series 1 and 2, with enumerated R-group sites where the free energies of various substituents were computed through alchemical calculations. (B) Comparison of experimentally measured free energy differences between substituents for series 1 (left) and 1 (right) for the best performing of the evaluated binding pose models. (C) Absolute binding free energies of all enantiomeric states of compounds A (left) and B (right) for the best performing pose of each ligand series.

The initial candidate poses were generated using the induced fit docking (IFD) protocol with extended sampling, as implemented in the Schödinger suite [25], which was run for five of the most potent ligands in each series in all their relevant enantiomeric states. For compounds in series 2, only endo-isomers were considered to faithfully replicate experimental conditions reported in [13], where exo-isomers were discarded. For each AlphaFold2 model, 400-500 candidate poses were generated, which we then characterized by generating their corresponding interaction fingerprints [26]. These fingerprints were clustered using a hierarchical algorithm, retaining 20 clusters per model based on cluster size and the top IFD score within each cluster.

Binding pose metadynamics [27] simulations were conducted to filter out unstable poses. Five replicas of 10 ns metadynamics were run per pose, evaluating stability through ligand RMSD and hydrogen bond conservation relative to the initial structure. The five most stable poses per model were selected for further evaluation.

To identify the pose most likely to represent the true bound state, we assessed how well the selected structures reproduced experimental binding affinities. A transformation network was constructed for 18 ligands in each series using the Lead Optimization Mapper [28]. Relative binding free energy (RBFE) differences were computed using alchemical transformations in OpenFE. Given that in the studied poses, the ligands are not exposed to the membrane or interact with lipid molecules at all, to streamline the analysis, we omitted the membrane in these RBFE and ABFE simulations. The predicted free energy differences were compared to experimental values using mean absolute error (MAE) and Kendall’s τ correlation, as shown in Figure 4B. For both series, at least one pose accurately reproduced experimental data, with MAE < 1.5 kcal/mol and τ > 0.4, indicating that these models capture key features of the true binding pose.

Beyond validating that these models could accurately predict relative free energy differences between ligands in the same series, we also assessed absolute binding free energies. ABFE calculations were performed for compounds A and B on the four top-scoring poses (based on correlation with experimental affinities), using the Yank program [29]. For both compounds, we identified at least one pose with a binding free energy closely matching experimental values, leading to the final top binding pose models (Figure 5A, 5B). For those selected models, the relative binding free energy calculations used to evaluate all candidate models were extended to include all ligands in each congeneric series. In both cases, we observed that the calculated free energy differences continued to closely match experimental data with MAE < 1.0 kcal/mol and τ > 0.4 in both cases.

**Figure 5.**
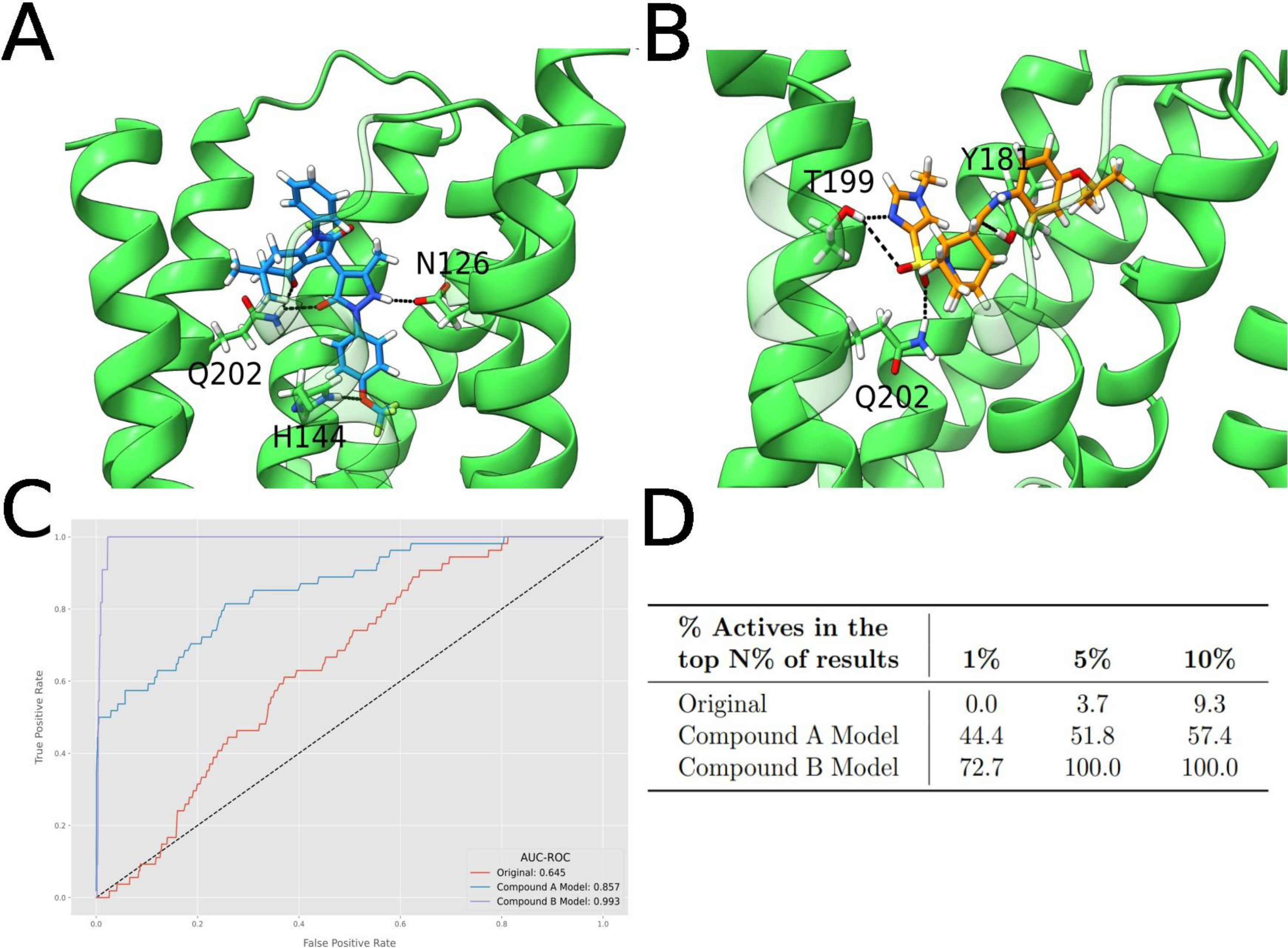
(A) Binding conformation of compound A in the optimal binding pose found for series 1. (B) Binding conformation of compound B in the optimal binding pose found for series 2. (C) ROC curves for the retrospective virtual screenings using the original AlphaFold model, and the optimal models for compounds A and B. (D) Percentage of active molecules recovered after selecting the top 1%, 5% and 10% ligands from the entire library, ranked by docking score.

To ensure the correct enantiomeric state was used in our models, we docked the alternative enantiomers with constraints to ensure that the resulting bound conformation was similar to the identified pose and ran ABFE calculations on all states. The results showed a preference for state S in compound A and state RRS in compound B (Figure 4C).

Analysis of the optimal bound poses of compounds A and B from both series 1 and 2, as depicted in Figures 5A and 5B, show that both occupy a similar volume in the binding pocket. Additionally, these bound poses are mediated by critical hydrogen bonds —ASP126 and HIS141 for compound A and THR199 for compound B, while both compounds also form hydrogen bonds with TYR181 and GLN202.

Trajectory analysis from the ABFE simulations indicated that most of these hydrogen bonds were stable over the majority of the simulation. Interestingly, although both TYR181 and GLN202 form hydrogen bonds with the initial docked poses of both compounds, simulations for compound A showed that the hydrogen bond with TYR181 is quickly lost during simulation, indicating that its role in stabilizing the bound poses may not be as important as GLN202, which maintained stable hydrogen bonding during the simulations of both compounds.

After identifying the optimal binding poses for each series, we re-evaluated the retrospective virtual screening analysis from section 2.2, to determine whether the conformations of these refined binding sites could provide better enrichments compared to the conformation of the binding site in the original AlphaFold model. As shown in figure 5C, both the series 1 and series 2 models showed significant improvements in terms or AUC-ROC compared to the original models. Importantly, these models did not show significant bias towards the ligand series for which they were optimized, as they both were able to correctly rank active ligands from the other series before most decoy molecules. Furthermore, we observed that constraining docking poses during the virtual screening analysis to form hydrogen bonds with GLN202 substantially improved the performance of all models (Figure 7 SI), and in particular increased the AUC-ROC of model A to match the performance of model B, further supporting the importance of this residue.

### Selectivity Analysis

Despite the structural similarity between the members of the ELOVL family, these enzymes are expressed in different tissues and have distinct fatty acid substrate preferences, which indicates that selectivity towards ELOVL6 is an important factor to consider when designing pharmacologically effective inhibitors. In addition to testing the activities of compounds A and B for ELOVL6, their selectivity was also experimentally validated by testing these compounds for members 1, 2, 3 and 5 of the ELOVL family [12, 13]. Compounds A and B showed 38 fold and 7 fold selectivity for ELOVL6 over ELOVL3 respectively, the member with the highest level of sequence homology to ELOVL6 (47%), and for which the second highest activity was measured. However, due to the lack of structural understanding of the inhibition mechanism, no understanding of the factors that induce selectivity in ELOVL enzymes exists yet.

To evaluate whether our binding mode models could address this knowledge gap, we adapted the ELOVL6 structural model to resemble other experimentally tested ELOVL family members. A sequence alignment identified non-conserved amino acids within 6 Å of the ligand in the predicted binding modes (Figure 6A). Keeping the binding poses of compounds A and B unchanged, we mutated these non-conserved residues to match those found in other ELOVL proteins. The resulting structures were energy-minimized using Prime, followed by ABFE calculations for both compounds. While this simplified approach may not fully capture all structural factors influencing selectivity, it allows us to determine whether direct interactions with non-conserved residues contribute significantly to ELOVL6 selectivity. If our models align with experimental selectivity trends, it suggests that interactions with specific non-conserved amino acids are key determinants of binding preference and should be considered when designing selective inhibitors.

**Figure 6.**
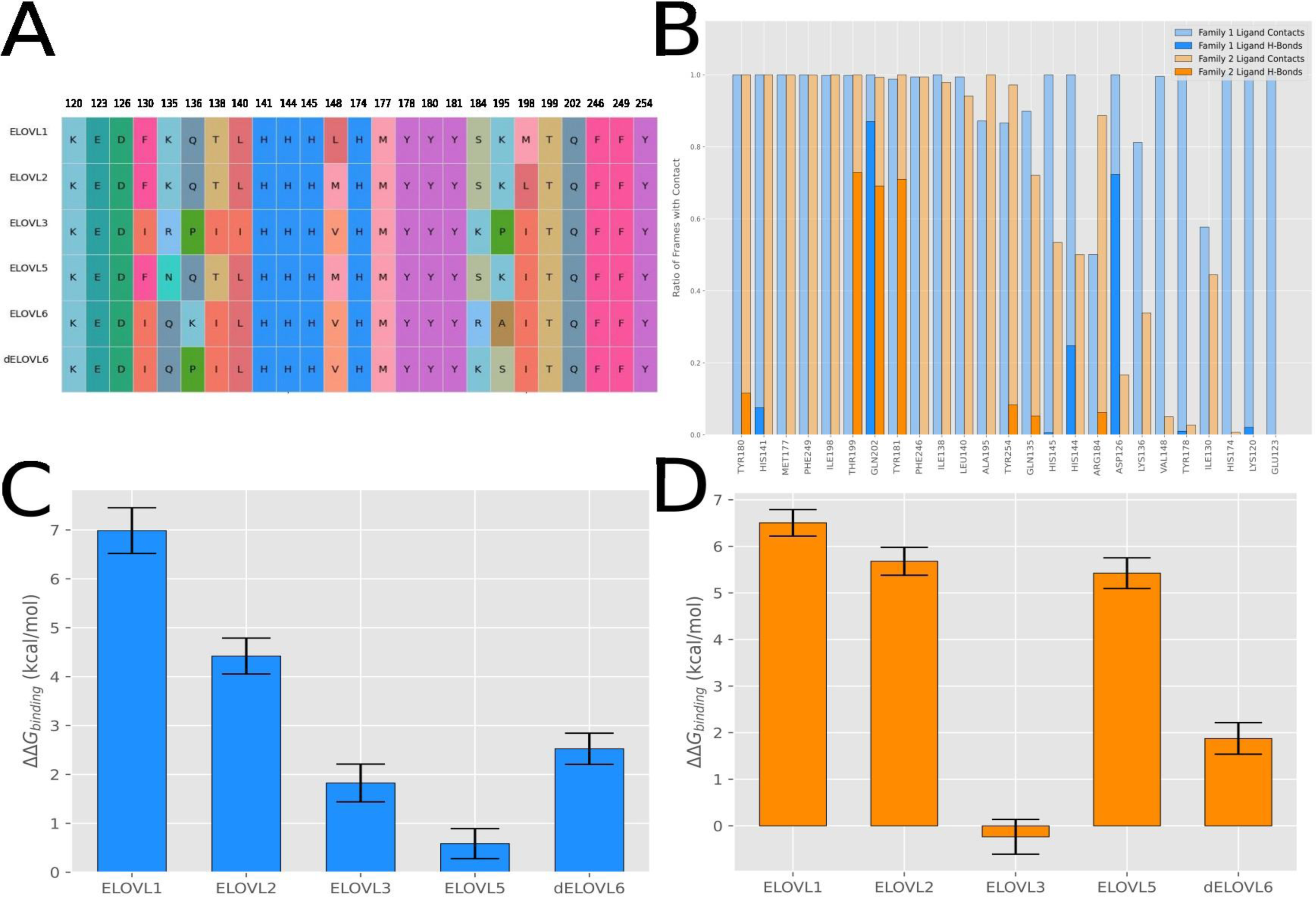
(A) Multiple sequence alignment of members 1, 2, 3, 5 and 6 of the ELOVL family, at the positions of ELOVL6 amino-acids that define the modeled binding pockets. (B) Contact frequency of the top 25 most commonly interacting residues in the ELOVL6 binding pocket for each compound. Bars represent the fraction of simulation frames in which a residue is in contact with the compound. Semi-transparent regions indicate general contacts, while opaque regions indicate hydrogen bond formation.. (C) Binding free energy differences between the bound pose of compound A in ELOVL6 and its bound pose in members 1, 2, 3 and 5 of the ELOVL family and member 6 of dELOVL, computed from ABFE calculations. (D) Binding free energy differences between the bound pose of compound B in ELOVL6 and its bound pose in members 1, 2, 3 and 5 of the ELOVL family and member 6 of dELOVL, computed from ABFE calculations.

For compound A, experimental results showed no measurable affinity for ELOVLs 1, 2 and 5, while IC_50_s for ELOVL6 and ELOVL3 were 8.9nm and 337nM, respectively [12]. Therefore, we would expect to see large differences, over 4kcal/mol, in the free energies of binding with respect to ELOVL6 for ELOVLs 1, 2 and 5, whereas ELOVL3 should have a binding free energy approximately 2kcal/mol lower than ELOVL6. A similar behavior would be expected for compound B, where members 1, 2 and 5 of the family showed no measurable affinity, whereas ELOVL3 showed a decreased affinity equivalent to approximately 2kcal/mol [13]. We also tested the affinity of these compounds for the ELOVL6 ortholog in *Drosophila melanogaster* (dELOVL6 or Baldspot), as the fruit fly is a highly accessible and suitable model for drug discovery in metabolic and neurological diseases. Furthermore, compound A has shown a significant effect in prolonging the lifespan of flies with amyotrophic lateral sclerosis pathology (Francisco J Gil-Bea, unpublished data), suggesting its inhibitory effects in ELOVL6 extend to dELOVL6.

Our ABFE calculations broadly reproduced these experimental trends (Figures 6C and 6D). For compound A, ELOVL3’s binding free energy was 1.8 kcal/mol lower than that of ELOVL6, while ELOVL1 and ELOVL2 exhibited differences exceeding 4 kcal/mol, consistent with experimental findings. For dELVOL6, the binding free energy was calculated to be 2.5 kcal/mol lower than for ELOVL6, which indicates that compound A should indeed have reasonable affinity for dELOVL6 as was experimentally measured. However, for ELOVL5, our model failed to capture significant free energy differences relative to ELOVL6, suggesting that structural differences beyond local binding site mutations may play a more prominent role in this case. Alternatively, our predicted binding pose may be missing key interactions that drive selectivity.

A similar pattern was observed for compound B, with the inactive ELOVL1, ELOVL2 and ELOVL5 models exhibiting lower binding affinities by 5 to 7 kcal/mol. Although no experimental verification of compound B’s affinity for dELOVL6 exists, our simulations suggest this compound may also be effective in *Drosophila*. However, unlike for compound A, for compound B our models were unable to capture the differences in affinity between ELOVL6 and ELOVL3, which reveals some limitations in our models for identifying the factors driving selectivity for these ligands.

Trajectory analysis of the ABFE simulations provided insights into the specific amino acids driving ELOVL6 selectivity. For ELOVL1, ELOVL2, and ELOVL5, multiple persistent ligand-contacting residues were mutated, particularly at positions 130, 148, 184 and 195, as well as in the 130–140 sequence region, which contains several non-conserved amino acids that persistently interacted with the inhibitor ligands during the 50ns ABFE simulations (Figure 6B). In contrast, ELOVL3 exhibits a more limited set of binding pocket mutations (GLN135, LYS136, LEU140, ARG184 and ALA195). Since LEU140 is mutated to the chemically similar isoleucine and ARG184 does not form stable interactions throughout the ABFE trajectory, we hypothesize that GLN135, LYS136 and ALA195 are the primary drivers of selectivity. Notably, the GLN135 and LYS136 mutations introduce charge changes, which could significantly alter the local electrostatic environment, contributing to the observed differences in binding affinity.

## DISCUSSION

In this work we have developed a binding mechanism model for ELOVL6 inhibitor molecules, and unveiled the key interactions that lead molecules to be strong binders, as well as the potentially crucial amino acids to consider when developing selective inhibitor molecules. Additionally, we developed structural models for ELOVL6, which demonstrated the ability to differentiate experimentally validated actives from decoy molecules with a high degree of accuracy. In fact, although these structural models were initially tailored to fit a specific congeneric series of molecules, their accuracy when differentiating active molecules with completely different chemical structures remained very high, indicating that this accuracy could be leveraged to find new lead compounds through virtual screening campaigns.

Although our understanding of the mechanism of action of ELOVL6 and its inhibition remains incomplete, we were able to provide insight into the likely binding pathway of FA substrates into ELOVL6. As far as the inhibition mechanism, previous studies [23] have suggested two main possible modes of action. The first is binding to an allosteric site of the enzyme’s active site, which could explain the selectivity of these ligands for ELOVL6 despite the similarity between ELOVL family members, and would additionally be consistent with the observation that these compounds inhibit palmitoyl-CoA uncompetitively, which could be explained by the fact that binding to this allosteric site might induce conformational changes that interfere with malonyl-CoA binding. We attempted to identify possible allosteric binding pockets in ELOVL6, but our models did not find allosteric sites that were able to reproduce experimental data for these inhibitors. Another possibility is that the inhibitor molecules recognize the conformational change associated with acyl-enzyme intermediate formation, and bind after this step in the catalytic process. Although a thorough analysis of the conformational changes that ELOVL6 goes through during the FA substrate elongation process is beyond the scope of this work, our results suggest that the affinity of these inhibitors for our modeled binding pockets is sensitive to slight realignments of the amino-acids in the binding pocket, as evidenced by the significant accuracy differences between the original AlphaFold model and our refined models in identifying active compounds.

Motivated by recent findings suggesting ELOVL6’s potential as a target for the treatment of various disorders spanning from metabolic, neurological and cancer malignancies [1–11], we hope to leverage these computational models to aid in the design of new ELOVL6 inhibitors with optimal pharmacokinetic properties in the future, both for lead identification through virtual screening of large-scale ligand libraries and lead optimization through RBFE calculations.

## ACKNOWLEDGMENTS

The authors gratefully acknowledge the financial support provided by the Departamento de Salud, Gobierno Vasco.

## REFERENCES

[1] Shimano, Hitoshi. “Novel qualitative aspects of tissue fatty acids related to metabolic regulation: lessons from Elovl6 knockout.” Progress in lipid research 51.3 (2012): 267–271.

[2] Matsuzaka, Takashi, et al. “Crucial role of a long-chain fatty acid elongase, Elovl6, in obesity-induced insulin resistance.” Nature medicine 13.10 (2007): 1193–1202.

[3] Matsuzaka, Takashi, et al. “Elovl6 promotes nonalcoholic steatohepatitis.” Hepatology 56.6 (2012): 2199–2208.

[4] Saito, Ryo, et al. “Macrophage Elovl6 deficiency ameliorates foam cell formation and reduces atherosclerosis in low-density lipoprotein receptor-deficient mice.” Arteriosclerosis, thrombosis, and vascular biology 31.9 (2011): 1973–1979.

[5] Guo, Jiao, et al. “Emerging insights on the role of Elovl6 in human diseases: Therapeutic challenges and opportunities.” Life Sciences (2024): 123308.

[6] Garcia Corrales, Aida V., et al. “Fatty acid elongation by ELOVL6 hampers remyelination by promoting inflammatory foam cell formation during demyelination.” Proceedings of the National Academy of Sciences 120.37 (2023): e2301030120.

[7] García García, Ana, et al. “Targeting ELOVL6 to Disrupt c-MYC Driven Lipid Metabolism in Pancreatic Cancer Enhances Chemosensitivity.” Nature Communications, vol. 16, no. 1, (2025), p. 1694.

[8] Muir, Kyle, et al. “Proteomic and lipidomic signatures of lipid metabolism in NASH-associated hepatocellular carcinoma.” Cancer research 73.15 (2013): 4722–4731.

[9] Anelli, Luisa, et al. “A novel t (4; 16)(q25; q23. 1) associated with EGF and ELOVL6 deregulation in acute myeloid leukemia.” Gene 529.1 (2013): 144–147.

[10] Shergalis, Andrea, et al. “Current challenges and opportunities in treating glioblastoma.” Pharmacological reviews 70.3 (2018): 412–445.

[11] Marien, Eyra, et al. “Phospholipid profiling identifies acyl chain elongation as a ubiquitous trait and potential target for the treatment of lung squamous cell carcinoma.” Oncotarget 7.11 (2016): 12582.

[12] Takahashi, Toshiyuki, et al. “Synthesis and evaluation of a novel indoledione class of long chain fatty acid elongase 6 (ELOVL6) inhibitors.” Journal of medicinal chemistry 52.10 (2009): 3142–3145.

[13] Sasaki, Takahide, et al. “Synthesis and evaluation of a novel 2-azabicyclo [2.2. 2] octane class of long chain fatty acid elongase 6 (ELOVL6) inhibitors.” Bioorganic & medicinal chemistry 17.15 (2009): 5639–5647.

[14] Nagase, Tsuyoshi, et al. “Synthesis and biological evaluation of a novel 3-sulfonyl-8-azabicyclo [3.2. 1] octane class of long chain fatty acid elongase 6 (ELOVL6) inhibitors.” Journal of medicinal chemistry 52.14 (2009): 4111–4114.

[15] Nie, Laiyin, et al. “The structural basis of fatty acid elongation by the ELOVL elongases.” Nature structural & molecular biology 28.6 (2021): 512–520.

[16] Jumper, John, et al. “Highly accurate protein structure prediction with AlphaFold.” Nature 596.7873 (2021): 583–589.

[17] Jacobson, Matthew P., et al. “A hierarchical approach to all-atom protein loop prediction.” Proteins: Structure, Function, and Bioinformatics 55.2 (2004): 351–367.

[18] Abraham, Mark James, et al. “GROMACS: High performance molecular simulations through multi-level parallelism from laptops to supercomputers.” SoftwareX 1 (2015): 19–25.

[19] Siddiqui, Arif Jamal, et al. “Computational insight into structural basis of human ELOVL1 inhibition.” Computers in Biology and Medicine 157 (2023): 106786.

[20] Bakan, Ahmet, et al. “Druggability assessment of allosteric proteins by dynamics simulations in the presence of probe molecules.” Journal of chemical theory and computation 8.7 (2012): 2435–2447.

[21] Cook, Charlie, et al. “Challenges of Absolute Binding Free Energies for Membrane Exposed Binding Pockets.” (2024).

[22] Ridsdale, Andrew, et al. “Cholesterol is required for efficient endoplasmic reticulum-to-Golgi transport of secretory membrane proteins.” Molecular biology of the cell 17.4 (2006): 1593–1605.

[23] Shimamura, Ken, et al. “5, 5-Dimethyl-3-(5-methyl-3-oxo-2-phenyl-2, 3-dihydro-1H-pyrazol-4-yl)-1-phenyl-3-(trifluoromethyl)-3, 5, 6, 7-tetrahydro-1H-indole-2, 4-dione, a potent inhibitor for mammalian elongase of long-chain fatty acids family 6: examination of its potential utility as a pharmacological tool.” Journal of Pharmacology and Experimental Therapeutics 330.1 (2009): 249–256.

[24] Mysinger, Michael M., et al. “Directory of useful decoys, enhanced (DUD-E): better ligands and decoys for better benchmarking.” Journal of medicinal chemistry 55.14 (2012): 6582–6594.

[25] Sherman, Woody, et al. “Novel procedure for modeling ligand/receptor induced fit effects.” Journal of medicinal chemistry 49.2 (2006): 534–553.

[26] Singh, Juswinder, et al. “Structural interaction fingerprints: a new approach to organizing, mining, analyzing, and designing protein–small molecule complexes.” Chemical Biology & Drug Design 67.1 (2006): 5–12.

[27] Fusani, Lucia, et al. “Exploring ligand stability in protein crystal structures using binding pose metadynamics.” Journal of Chemical Information and Modeling 60.3 (2020): 1528–1539.

[28] Liu, Shuai, et al. “Lead optimization mapper: automating free energy calculations for lead optimization.” Journal of computer-aided molecular design 27 (2013): 755–770.

[29] Rizzi A, Grinaway PB, Parton DL, Shirts MR, Wang K, Eastman P, Friedrichs M, Pande VS, Branson K, Mobley DL, Chodera JD. YANK: A GPU-accelerated platform for alchemical free energy calculations. In preparation.

