## Supplementary Material for "Computational Analysis of ELOVL6 Structure and Inhibition for Rational Drug Design"

### Molecular Dynamics Simulation Details

#### Umbrella Sampling Simulations

To generate initial configurations for umbrella sampling, steered molecular dynamics (SMD) simulations were performed to explore both hypothesized substrate binding mechanisms: lateral and sequential insertion. Ten independent SMD replicas were initiated from the same starting structure. In each replica, a moving harmonic restraint was applied to the ligand to steer it along the proposed binding coordinate. The pulling rate was set to 0.01 nm/ps, and a force constant of 1000 kJ $mol^{-1}$ $nm^{-2}$ was used for the moving harmonic potential. The total work performed during each trajectory was computed, and the three trajectories with the lowest total work were selected for further analysis.

From each selected trajectory, 25 structures were extracted at equidistant intervals along the reaction coordinate to define the umbrella sampling windows. A separate umbrella sampling simulation was carried out for each of the three structure sets, resulting in three umbrella sampling replicates.

All simulations were performed using GROMACS 2024.1 [1] with the amber ff14SB force field for the protein [2] and GAFF2 for the substrate [3]. Systems were solvated in TIP3P water [4] with 0.15 M KCl, and long-range electrostatics were treated using Particle Mesh Ewald (PME) [5]. Short-range interactions were truncated at 9 Å. Temperature was maintained at 300 K using Langevin dynamics, and pressure was kept at 1 atm using the stochastic cell rescaling barostat [6].

Each SMD replica underwent energy minimization, followed by 0.5 ns of NVT and 0.5 ns of NPT equilibration, with positional restraints applied to protein heavy atoms. Production SMD simulations were run for 1 ns with a 2 fs timestep, applying the moving harmonic potential described above.

For each of the three selected SMD trajectories, 25 umbrella sampling windows were defined using the extracted structures. Each window was equilibrated for 10 ns under NPT conditions, followed by 75 ns of production simulation. A fixed harmonic bias with a force constant of 1000 kJ $mol^{-1}$ $nm^{-2}$ was applied in each window to restrain the system along the reaction coordinate. Simulations were conducted using the same thermostat, barostat, and cutoff settings described above, with a 2 fs timestep.

Free energy profiles along the reaction coordinate were computed using the WHAM implementation provided in GROMACS 2024.1.

#### Mixed Solvent Molecular Dynamics

Mixed solvent MD simulations were performed to identify potential ligand interaction hotspots on the protein surface. Five independent replicas were run, with simulation times summarized in Table 1.

All systems were built following the protocol by Bakan et al. [1]. After energy minimization, a 0.6 ns annealing protocol was applied: the system was heated to 500 K and cooled to 300 K while restraining the heavy atoms of the protein. This was followed by 0.6 ns of NPT equilibration.

Simulations were run using NAMD 3.0 [2] with the CHARMM36m force field [3]. The systems were solvated with TIP3P water [4], and $Cl^-$ ions were added for charge neutrality. Probe molecules (isopropanol, acetamide, acetate, isopropylamine, and benzene) were included at 2 M concentration.

A 2 fs timestep was used with a 12 Å cutoff for nonbonded interactions, and long-range electrostatics were treated using Particle Mesh Ewald (PME) [5]. Temperature was maintained at 300 K via Langevin dynamics, and pressure was kept at 1 atm using the Langevin piston method.

Spatial distribution functions of probe molecules were analyzed using DruGUI for VMD [6], and volumetric maps were visualized in ChimeraX [7].

| | Replica 1 | Replica 2 | Replica 3 | Replica 4 | Replica 5 |
|---|---|---|---|---|---|
| Simulation Time (ns) | 267 | 274 | 273 | 268 | 274 |

**Absolute Binding Free Energy (ABFE) with Membrane**

ABFE simulations incorporating the presence of an explicit membrane were conducted using GROMACS 2024.1 [8]. The systems consisted of membrane-embedded protein-ligand complexes, built via CHARMM-GUI [9] using a lipid bilayer of 90% POPC and 10% cholesterol.

The amber ff14SB [10], GAFF2 [11], and LIPID21 [12] force fields were used for the protein, ligand, and lipids, respectively. Solvation was performed with TIP3P water and 0.15 M KCl.

Each λ-window underwent energy minimization, followed by 0.5 ns of NPT equilibration. The production phase involved 40 ns simulation for the complex phase and 10 ns for the solvent phase. A 2 fs timestep was used with a 9 Å cutoff for short-range interactions and PME for long-range electrostatics.

Simulations were performed at 300 K using Langevin dynamics, and pressure was maintained at 1 atm using a semi-isotropic stochastic cell rescaling barostat [13] in the complex phase.

Boresch restraints [14] were applied to fix the bound ligand conformation, with atoms selected using MDRestraintsGenerator [15]. A total of 42 λ-windows were used in the complex phase and 21 in the solvent phase. Free energy differences were calculated using the BAR method [16] as implemented in GROMACS 2024.1.

**Absolute Binding Free Energy (ABFE) in Solution**

ABFE simulations in aqueous solution were performed using Yank v0.25.2 [17], with OpenMM 8.1.1 [18] as the simulation engine. The amber ff14SB [10] and GAFF2 [11] force fields were used for the protein and ligands, respectively. The systems were solvated with TIP3P water and 0.15 M KCl.

Each λ-window underwent energy minimization and 1 ns equilibration, followed by 40 ns of production simulation for both the complex and solvent phases. Hamiltonian replica exchange [19] was used between windows to enhance sampling.

Simulations used a 4 fs timestep, Langevin dynamics at 300 K, and a Monte Carlo barostat [20] at 1 atm. Short-range interactions were truncated at 9 Å, and long-range electrostatics were computed using PME [5].

Boresch restraints were applied automatically by Yank. 42 λ-windows were used for the complex phase and 29 for the solvent phase. Final free energy estimates were obtained using

Yank's internal analysis tools. Contact and hydrogen bond analyses were performed with MDTraj [21] and MDAnalysis [22].

**Relative Binding Free Energy (RBFE)**

RBFE simulations were performed using OpenFE v0.25.2 [23], with OpenMM 8.1.1 [18] as the simulation engine. The amber ff14SB [10] and GAFF2 [11] force fields were used for proteins and ligands, respectively. All systems were solvated in TIP3P water with 0.15 M KCl.

Each λ-window was subjected to energy minimization and 1 ns NPT equilibration, followed by 5 ns of production simulation for both phases. For transformations showing poor convergence, simulation time was extended to 20 ns.

Simulations were carried out using Hamiltonian replica exchange [19] across 11 λ-windows, with expansion to 25 windows where needed due to poor overlap between adjacent windows in the MBAR matrix. The temperature was kept at 300 K using Langevin dynamics, and pressure was held at 1 atm using a Monte Carlo barostat [20]. A 4 fs timestep and 9 Å cutoff were used. PME was used for long-range electrostatics.

For the evaluation of models generated through IFD, a single replica per transformation was used, while 3 replicas per transformation were used for the validation of the best performing models. The ligand transformation network was computed based on Lomap scores [24], and absolute binding free energies were derived from relative binding free energies using the method implemented in [25].

Table 1: Docking scores, induced fit docking scores and CompScore from binding pose metadynamics simulations, computed following [], for compound A. Data for each of the 5 best selected clusters obtained from the IFD simulations of all 5 AlphaFold models is shown.

| **Model** | **Cluster** | **CompScore** | **IFDScore** | **Docking Score** |
|---|---|---|---|---|
| Model 1 | Cluster 1 | -3.48 | -11376.91 | -11.96 |
| | Cluster 2 | 1.45 | -11330.38 | -12.38 |
| | Cluster 3 | -1.44 | -11285.65 | -8.90 |
| | Cluster 4 | -2.35 | -11338.03 | -9.89 |
| | Cluster 5 | 1.72 | -11313.85 | -8.73 |
| Model 2 | Cluster 1 | -2.17 | -11319.64 | -11.09 |
| | Cluster 2 | -1.21 | -11300.67 | -8.08 |
| | Cluster 3 | -0.78 | -11313.93 | -10.49 |
| | Cluster 4 | 1.33 | -11331.27 | -11.89 |
| | Cluster 5 | -1.35 | -11313.67 | -10.04 |
| Model 3 | Cluster 1 | -1.05 | -11345.90 | -10.00 |
| | Cluster 2 | 0.21 | -11362.96 | -11.21 |
| | Cluster 3 | -0.67 | -11312.65 | -9.02 |
| | Cluster 4 | -0.08 | -11408.97 | -11.32 |
| | Cluster 5 | -0.71 | -11306.65 | -10.22 |
| Model 4 | Cluster 1 | -0.36 | -11391.44 | -9.11 |
| | Cluster 2 | 0.43 | -11399.33 | -10.15 |
| | Cluster 3 | 1.10 | -11357.53 | -10.41 |
| | Cluster 4 | -0.59 | -11386.28 | -9.15 |
| | Cluster 5 | -1.88 | -11387.47 | -10.60 |
| Model 5 | Cluster 1 | -0.07 | -11303.85 | -10.32 |
| | Cluster 2 | -1.13 | -11372.21 | -9.93 |
| | Cluster 3 | -1.20 | -11360.27 | -9.37 |
| | Cluster 4 | -0.20 | -11304.95 | -11.26 |
| | Cluster 5 | 1.63 | -11337.46 | -11.08 |

Table 2: Docking scores, induced fit docking scores and CompScore from binding pose metadynamics simulations, computed following [], for compound B. Data for each of the 5 best selected clusters obtained from the IFD simulations of all 5 AlphaFold models is shown.

| Model | Cluster | CompScore | IFDScore | Docking Score |
|---|---|---|---|---|
| Model 1 | Cluster 1 | -1.03 | -11384.68 | -8.45 |
| | Cluster 2 | 1.54 | -11414.41 | -8.70 |
| | Cluster 3 | -0.15 | -11417.89 | -9.11 |
| | Cluster 4 | -0.53 | -11373.11 | -9.95 |
| | Cluster 5 | 0.02 | -11381.47 | -10.36 |
| Model 2 | Cluster 1 | -0.93 | -11326.63 | -9.00 |
| | Cluster 2 | 1.28 | -11323.93 | -7.51 |
| | Cluster 3 | 1.28 | -11306.00 | -7.16 |
| | Cluster 4 | 0.43 | -11290.96 | -7.35 |
| | Cluster 5 | -1.83 | -11357.01 | -9.03 |
| Model 3 | Cluster 1 | -0.08 | -11341.53 | -7.40 |
| | Cluster 2 | -0.78 | -11368.36 | -9.21 |
| | Cluster 3 | 1.68 | -11377.64 | -9.22 |
| | Cluster 4 | -2.23 | -11384.47 | -9.20 |
| | Cluster 5 | 0.25 | -11338.72 | -8.12 |
| Model 4 | Cluster 1 | 2.44 | -11386.66 | -9.07 |
| | Cluster 2 | -0.68 | -11370.32 | -9.55 |
| | Cluster 3 | -0.21 | -11389.09 | -7.19 |
| | Cluster 4 | 1.93 | -11403.79 | -8.47 |
| | Cluster 5 | -1.88 | -11302.75 | -6.86 |
| Model 5 | Cluster 1 | 0.06 | -11396.06 | -9.53 |
| | Cluster 2 | -2.99 | -11386.91 | -7.24 |
| | Cluster 3 | 0.45 | -11416.65 | -8.00 |
| | Cluster 4 | -1.22 | -11403.45 | -7.10 |
| | Cluster 5 | 2.06 | -11391.49 | -8.21 |

**Figure 1**. Backbone RMSD values of the best AlphaFold model for ELOVL6 and the crystal structure of ELOVL7 (PDBID 6Y7F) obtained from independent simulations of both proteins at temperature of 300K, 400K and 500K.

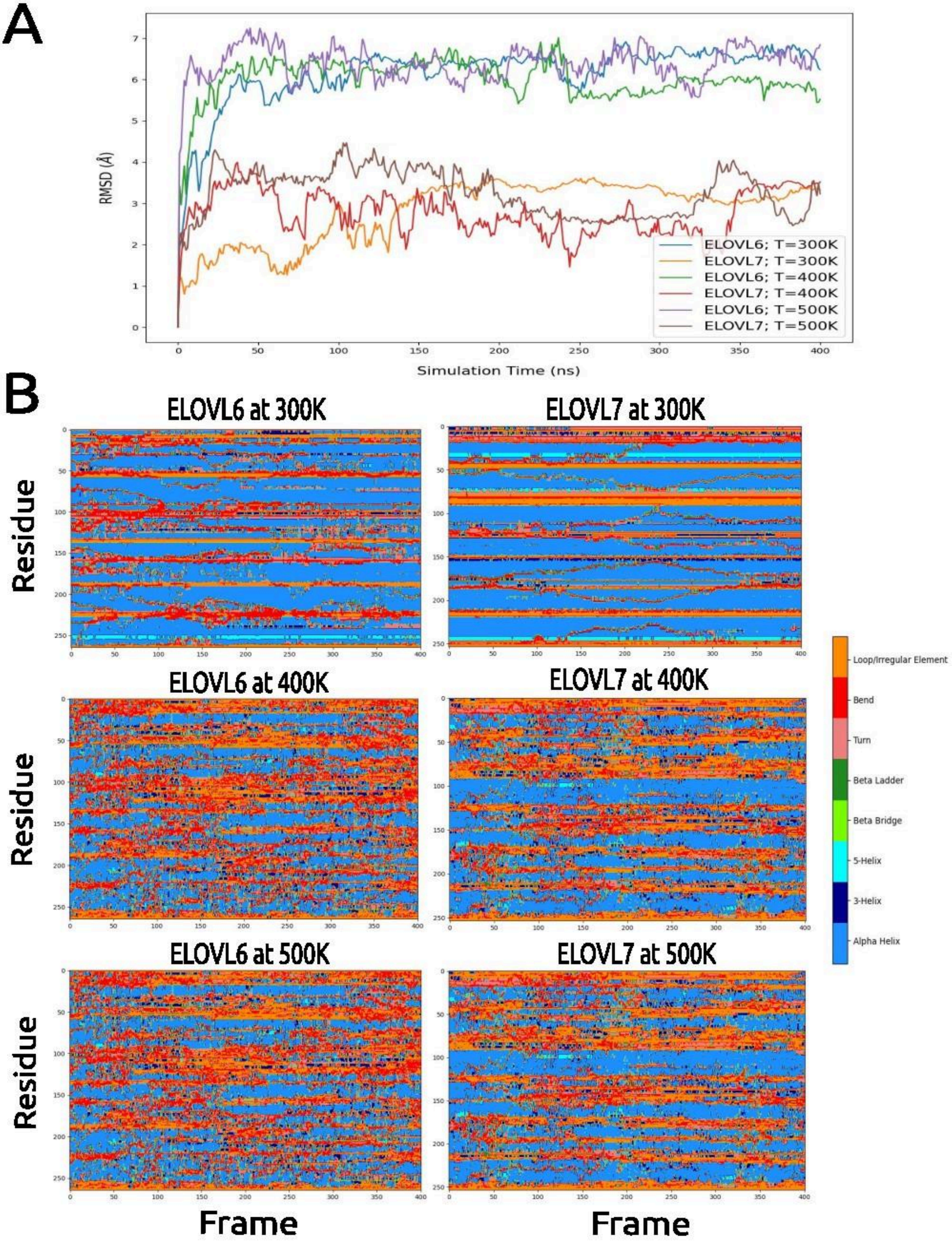

**Figure 2**. Schematic representation of the protocol used for the generation and validation of bound poses based on their consistency with experimental data.

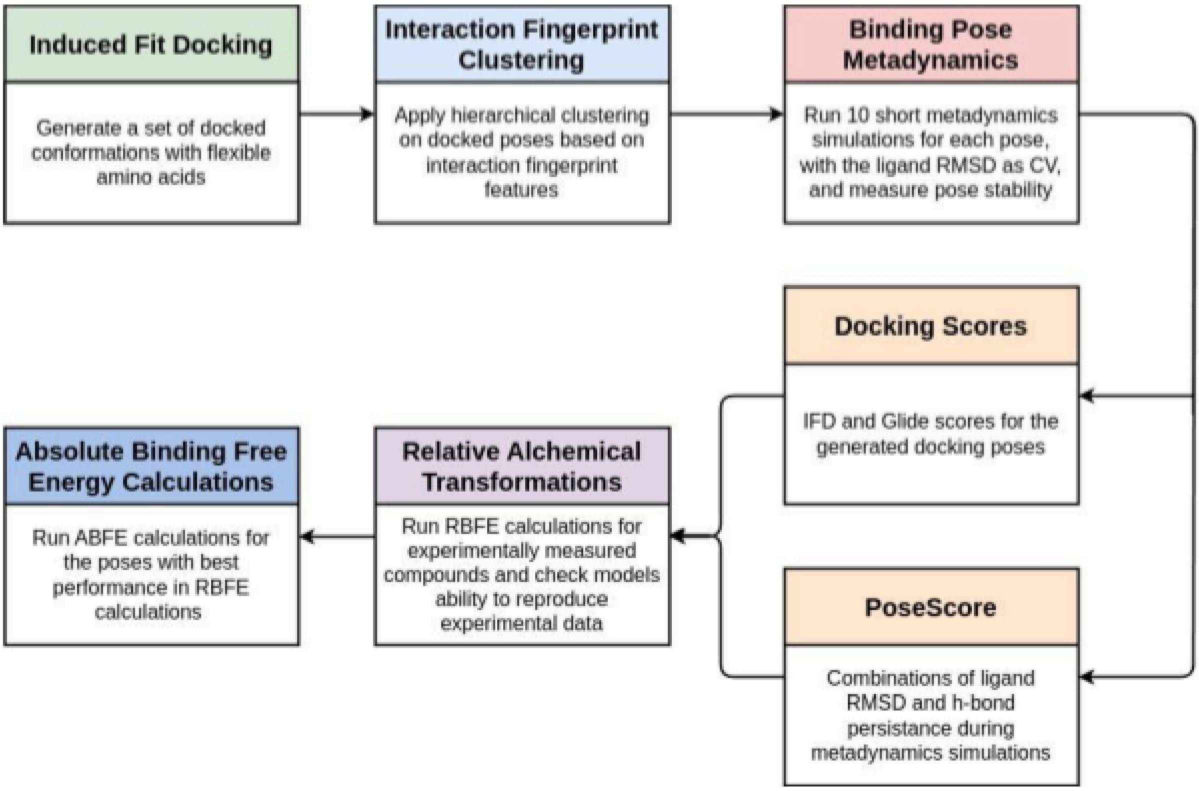


**Figure 3**. Heatmap representing the performance of all bound conformations of compound A, evaluated through the correlation between experimental binding free energy differences and binding free energy differences calculated from RBFE simulations of a subset of compound A's derivatives.

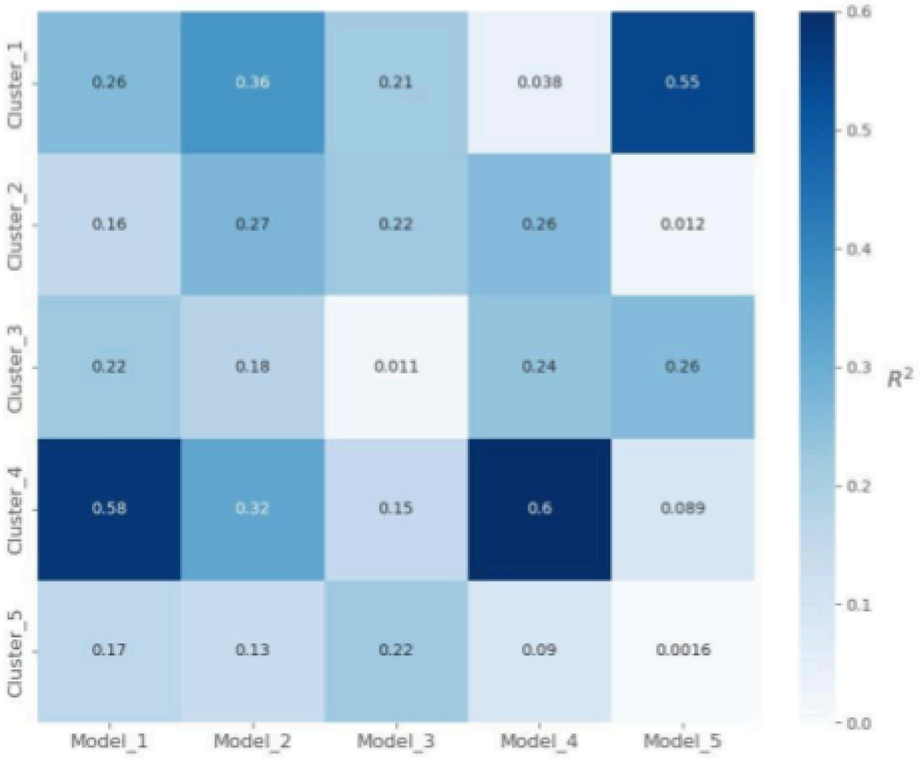

**Figure 4**. Heatmap representing the performance of all bound conformations of compound B, evaluated through the correlation between experimental binding free energy differences and binding free energy differences calculated from RBFE simulations of a subset of compound B's derivatives.

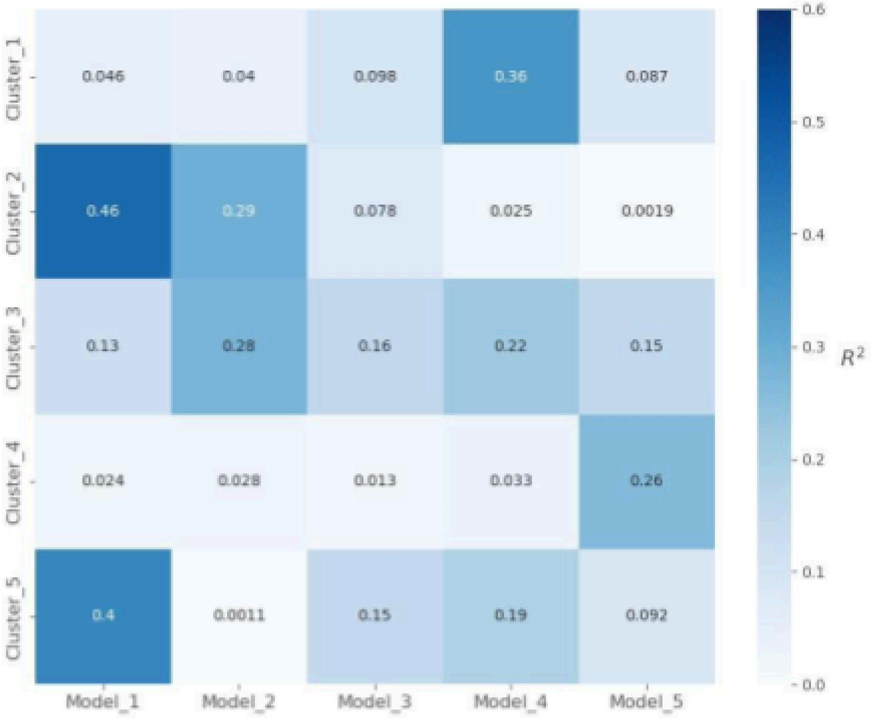


**Figure 5**. Absolute binding free energies for the 3 best performing IFD poses of compound A, based on the correlation values from figure 3. Poses with coefficients of determination above 0.4 were simulated.

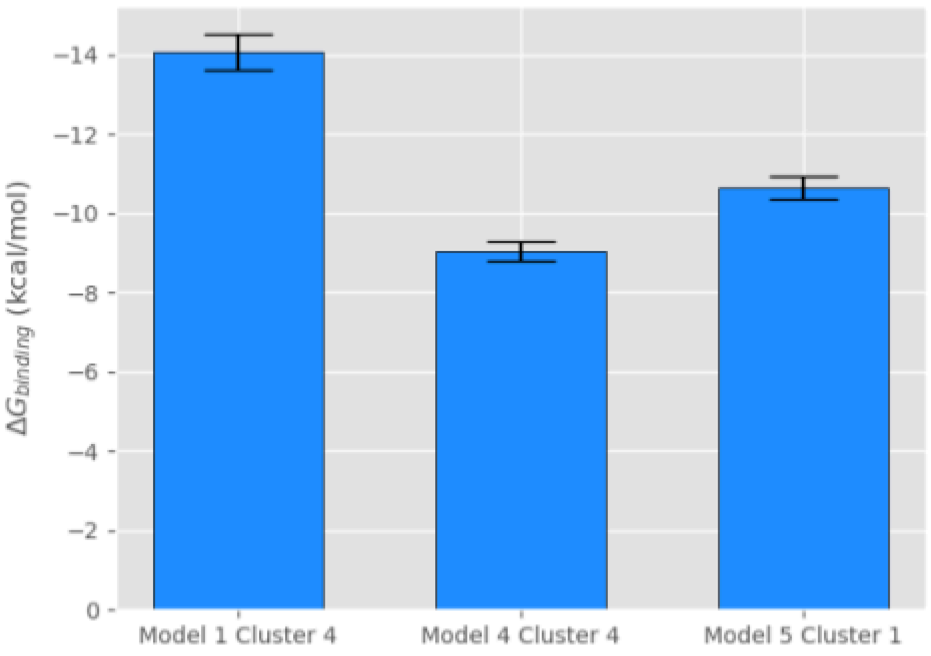

**Figure 6**. Absolute binding free energies for the 2 best performing IFD poses of compound B, based on the correlation values from figure 4. Poses with coefficients of determination above 0.4 were simulated.

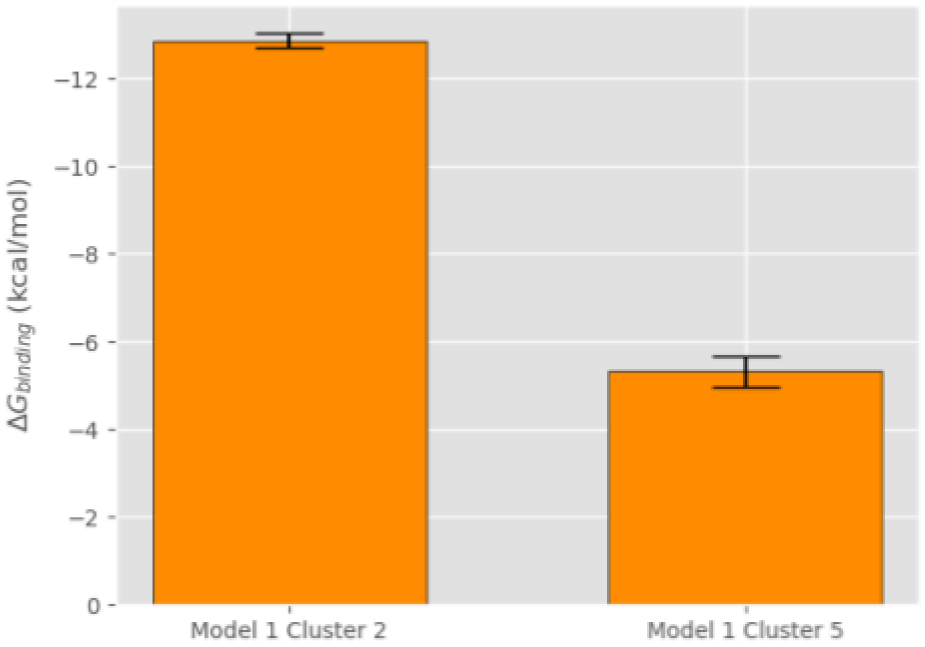


**Figure 7**. ROC curves for the retrospective virtual screenings using the original AlphaFold model, and the optimal models for ligand series 1 and 2, with constraints on the allowed docking poses to require hydrogen bond formation with residues TYR181 and/or GLN202.

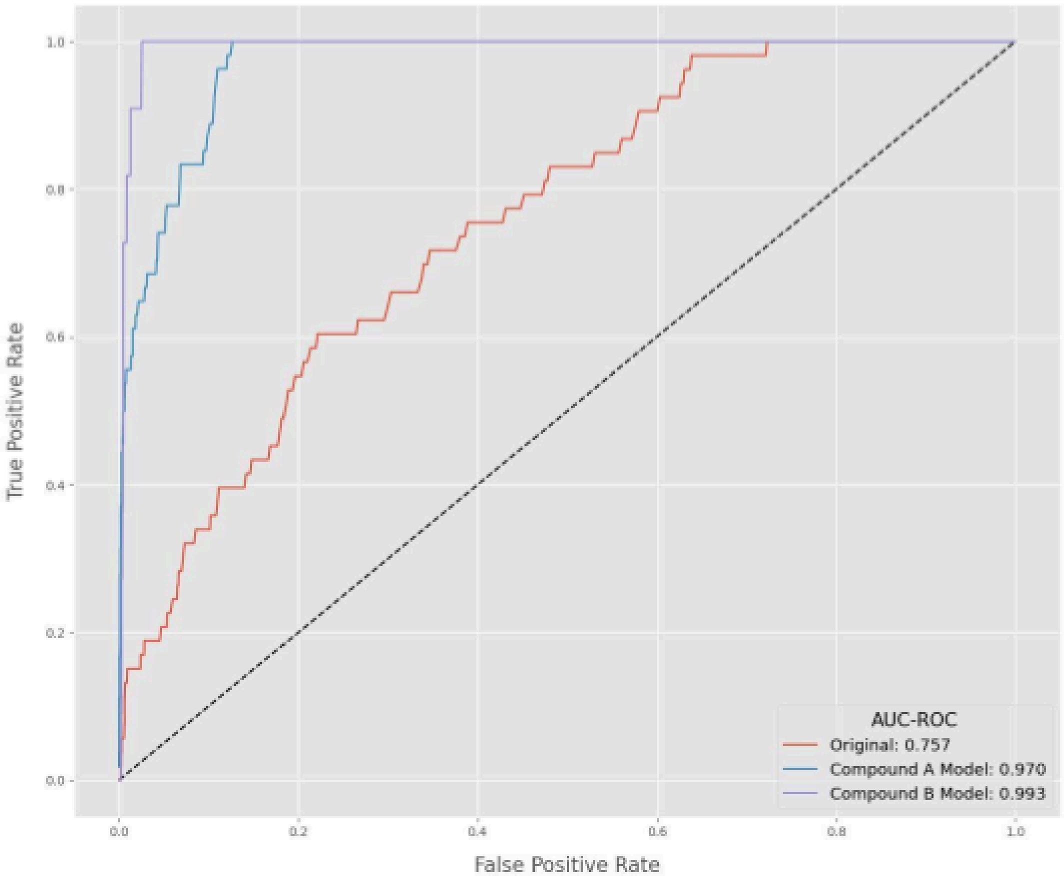

**Figure 8**. Comparison between the the docking score distributions of known active molecules and decoy molecules for (**A**) the original AlphaFold model, (**B**) optimal model for compound A and (**C**) optimal model for compound B, showing a clear distributional shift towards better docking scores for known actives compared to decoys.

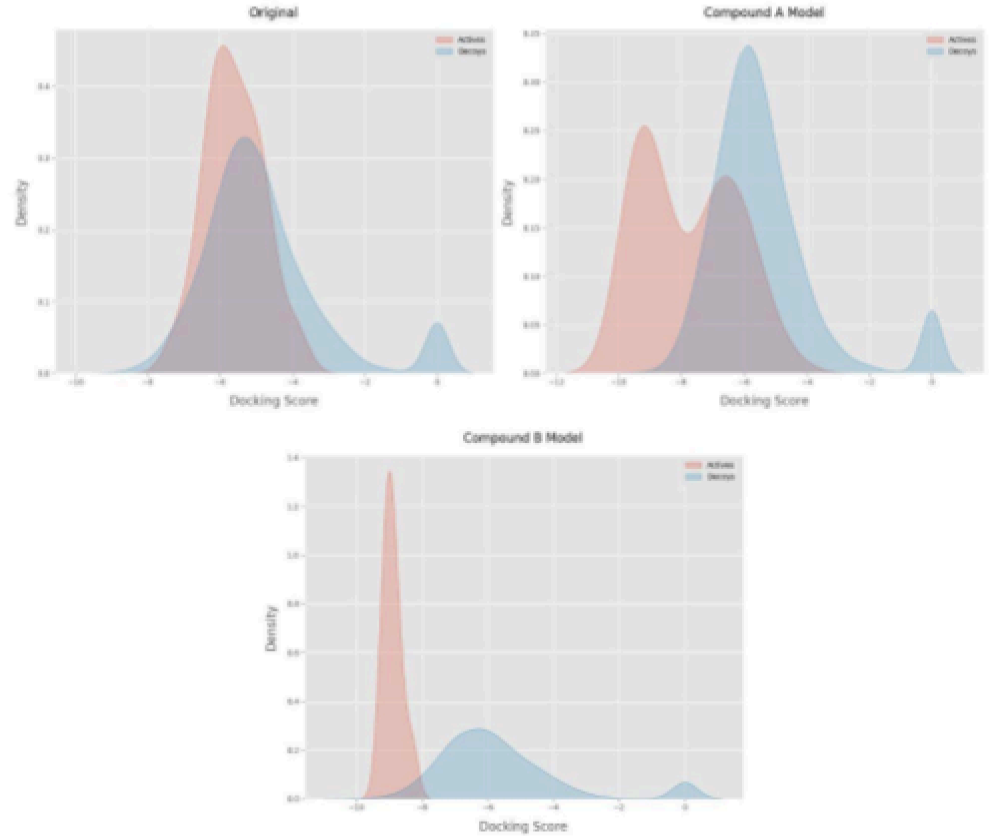


#### References


1. Bakan, A. et al. Druggability assessment of allosteric proteins by dynamics simulations in the presence of probe molecules. *J. Chem. Theory Comput.* 2012, *8*, 2435–2447.

2. Phillips, J. C. et al. Scalable molecular dynamics on CPU and GPU architectures with NAMD. *J. Chem. Phys.* 2020, *153*, 044130.

3. Huang, J.; MacKerell, A. D. CHARMM36m: An improved force field for folded and intrinsically disordered proteins. *J. Comput. Chem.* 2017, *34*, 2135–2145.

4. Jorgensen, W. L. et al. Comparison of simple potential functions for simulating liquid water. *J. Chem. Phys.* 1983, *79*, 926.

5. Darden, T. et al. Particle mesh Ewald: An N·log(N) method for Ewald sums in large systems. *J. Chem. Phys.* 1993, *98*, 10089–10092.

6. Humphrey, W. et al. VMD: Visual molecular dynamics. *J. Mol. Graph.* 1996, *14*, 33–38.

7. Pettersen, E. F. et al. UCSF ChimeraX: Structure visualization for researchers, educators, and developers. *Protein Sci.* 2021, *30*, 70–82.

8. Abraham, M. J. et al. GROMACS: High performance molecular simulations. *SoftwareX* 2015, *1–2*, 19–25.

9. Jo, S. et al. CHARMM-GUI: A web-based graphical user interface for CHARMM. *J. Comput. Chem.* 2008, *29*, 1859–1865.

10. Maier, J. A. et al. ff14SB: Improving the accuracy of protein side chain and backbone parameters. *J. Chem. Theory Comput.* 2015, *11*, 3696–3713.

11. Wang, J. et al. Development and testing of a general amber force field. *J. Comput. Chem.* 2004, *25*, 1157–1174.

12. Hsu, P. C. et al. LIPID21: A comprehensive force field for cholesterol and phospholipids. *J. Chem. Theory Comput.* 2022, *18*, 6948–6960.

13. Bussi, G. et al. Isothermal-isobaric molecular dynamics using stochastic velocity rescaling. *J. Chem. Phys.* 2009, *130*, 074101.

14. Boresch, S. et al. Absolute binding free energies: A quantitative approach for their calculation. *J. Phys. Chem. B* 2003, *107*, 9535–9551.

15. Hahn, D. F. et al. MDRestraintsGenerator: Restraint generation for free energy calculations. *J. Chem. Inf. Model.* 2023, *63*, 1133–1141.

16. Bennett, C. H. Efficient estimation of free energy differences from Monte Carlo data. *J. Comput. Phys.* 1976, *22*, 245–268.

17. Chodera, J. D. et al. Yank: GPU-accelerated platform for alchemical free energy calculations. *J. Chem. Theory Comput.* 2018, *14*, 6343–6354.

18. Eastman, P. et al. OpenMM 7: Rapid development of high performance molecular dynamics algorithms. *PLoS Comput. Biol.* 2017, *13*, e1005659.

19. Sugita, Y.; Okamoto, Y. Replica-exchange molecular dynamics for protein folding. *Chem. Phys. Lett.* 1999, *314*, 141–151.

20. Shirts, M. R.; Chodera, J. D. Statistically optimal analysis of samples from multiple equilibrium states. *J. Chem. Phys.* 2008, *129*, 124105.

21. McGibbon, R. T. et al. MDTraj: A modern open library for the analysis of molecular dynamics trajectories. *Biophys. J.* 2015, *109*, 1528–1532.

22. Michaud-Agrawal, N. et al. MDAnalysis: A toolkit for the analysis of molecular dynamics simulations. *J. Comput. Chem.* 2011, *32*, 2319–2327.

23. OpenFE Development Team. OpenFE: Open-source toolkit for relative binding free energy calculations. https://github.com/OpenFreeEnergy
24. Liu, Shuai, et al. "Lead optimization mapper: automating free energy calculations for lead optimization." *Journal of computer-aided molecular design* 27 (2013): 755-770.
25. Li, Yishui, et al. "An open source graph-based weighted cycle closure method for relative binding free energy calculations." *Journal of Chemical Information and Modeling* 63.2 (2022): 561-570.
